# Widespread changes in gene expression accompany body size evolution in nematodes

**DOI:** 10.1101/2023.10.30.564729

**Authors:** Gavin C. Woodruff, John H. Willis, Erik Johnson, Patrick C. Phillips

## Abstract

Body size is a fundamental trait that drives multiple evolutionary and ecological patterns. *Caenorhabditis inopinata* is a fig-associated nematode that is exceptionally large relative to other members of the genus, including *C. elegans*. We previously showed that *C. inopinata* is large primarily due to postembryonic cell size expansion that occurs during the larval-to-adult transition. Here, we describe gene expression patterns in *C. elegans* and *C. inopinata* throughout this developmental period to understand the transcriptional basis of body size change. We performed RNA-seq in both species across the L3, L4, and adult stages. Most genes are differentially expressed across all developmental stages, consistent with *C. inopinata*’s divergent ecology and morphology. We also used a model comparison approach to identify orthologs with divergent dynamics across this developmental period between the two species. This included genes connected to neurons, behavior, stress response, developmental timing, and small RNA/chromatin regulation. Multiple hypodermal collagens were also observed to harbor divergent developmental dynamics across this period, and genes important for molting and body morphology were also detected. Genes associated with TGF-β signaling revealed idiosyncratic and unexpected transcriptional patterns given their role in body size regulation in *C. elegans*. Widespread transcriptional divergence between these species is unexpected and may be a signature of the ecological and morphological divergence of *C. inopinata*. Alternatively, transcriptional turnover may be the rule in the *Caenorhabditis* genus, indicative of widespread developmental system drift among species. This work lays the foundation for future functional genetic studies interrogating the bases of body size evolution in this group.

## Introduction

The size of an organism is both conspicuous and central to its way of life. Life history strategies are intimately tied to body size; for instance, as larger organisms tend to develop more slowly (McMahon and Bonner 1983; Calder 1984), body size underlies trade-offs between maturation time and larval survival (Stearns 1992). Body size dictates the kinds of organisms one directly interacts with as well as the nature of those interactions (Peters 1983; Calder 1984). Moreover, the physical space an organism occupies dictates the scale of its influence on the environment (*i*.*e*., body size correlates with home range: a single bacterial cell’s spatial sphere of influence is vastly different from that of a single blue whale) (Peters 1983; Calder 1984). As a consequence, the diversity of body sizes in the natural world is immense (twenty-one orders of magnitude (McMahon and Bonner 1983)) and obvious. A satisfying explanation of diversity will then require an account of the causes (both proximate and ultimate) of body size diversity.

One approach towards understanding how body sizes change is to study closely-related organisms with divergent body sizes where the genetic traces of the bases of evolutionary change are still detectable. The nematode genus *Caenorhabditis* is well-positioned to address this problem— *C. elegans* is a model system with sophisticated genetic tools (Corsi *et al*. 2015), and its sister species, *C. inopinata*, has rapidly evolved a much larger body size (being 64-200% longer in body length (Woodruff *et al*. 2018; Kanzaki *et al*. 2018)). In a previous study, we showed that this body size difference largely occurs due to postembryonic events during the larval to adult transition (Woodruff *et al*. 2018) (Fig. 1). Additionally, we showed that this difference was not due to changes in cell number nor epidermal ploidy (Woodruff *et al*. 2018). We then concluded that changes in cell size upon maturation was the major driver of body size divergence in this system (Woodruff *et al*. 2018).

**Figure 1.**
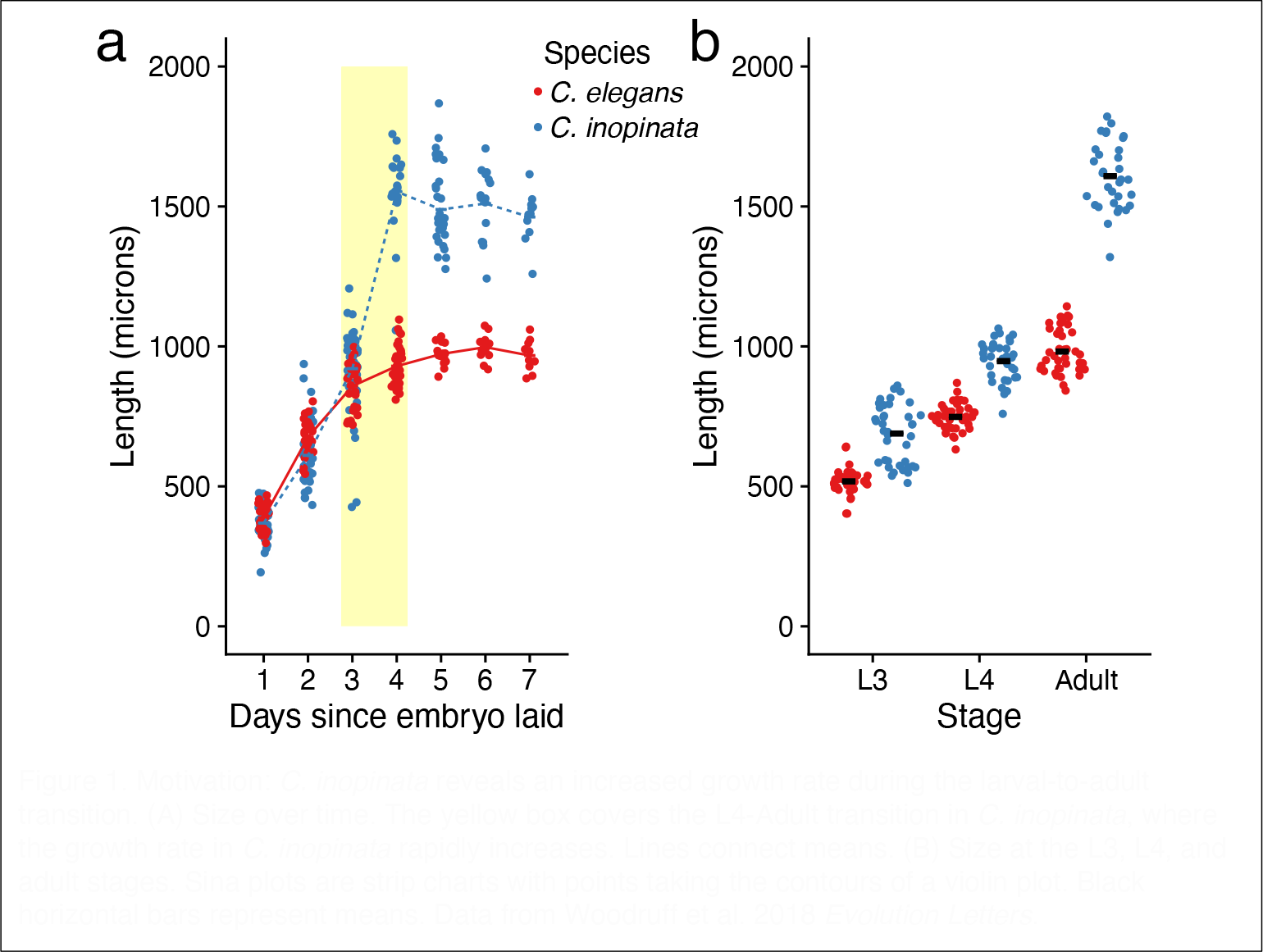
Motivation: *C. inopinata* reveals an increased growth rate during the larval-to-adult transition. (A) Size over time. The yellow box covers the L4-Adult transition in *C. inopinata*, where the growth rate in *C. inopinata* rapidly increases. Lines connect means. (B) Size at the L3, L4, and adult stages. Sina plots are strip charts with points taking the contours of a violin plot. Black horizontal bars represent means. Data from Woodruff et al. 2018 *Evolution Letters*.

Another advantage of the *Caenorhabditis* system is the vast body of background knowledge associated with *C. elegans* (Corsi *et al*. 2015). A number of body size mutants have been isolated in this system, providing a wealth of genetic fodder for evolutionary hypotheses regarding the bases of body size divergence in *C. inopinata*. For instance, multiple genes in the Transforming Growth Factor β (TGF-β) pathway reveal body size phenotypes when perturbed with mutation or RNAi (Savage-Dunn and Padgett 2017). Moreover, there are dose-dependent responses of TGF-β signaling factors on body size in *C. elegans* (Suzuki *et al*. 1999; Morita *et al*. 1999). Thus, one hypothesis for the evolution of body size in *C. inopinata* would be the modulation of TGF-β pathway activity via changes in gene expression. Likewise, multiple cuticle collagens also regulate body size in *C. elegans* (Johnstone 2000; Madaan *et al*. 2018); changes in their copy number or expression could also underlie body size evolution in *C. inopinata*. Indeed, hundreds of genes in *C. elegans* have been shown to influence body size (Schindelman *et al*. 2011), and all of them represent hypothetical drivers of body size divergence in *C. inopinata*. To test these hypotheses, we performed RNA-seq in *C. elegans* and *C. inopinata* across the L3-adult transition to find genes with divergent developmental dynamics that are potentially connected to the evolution of body size.

## Materials & Methods

### Strains, culture conditions, and developmental staging

*C. inopinata* NKZ35 (Kanzaki *et al*. 2018) and *C. elegans fog-2(q71)* JK574 (Schedl and Kimble 1988) were used for this study. Animals were maintained on Nematode Growth Media (with 3.2% agar) seeded with the food source *Escherichia coli* OP50-1 at 25°C. Animals were synchronized by allowing 15 adult *C. elegans* (or 70 adult *C. inopinata*) gravid females to lay embryos for three hours. 15 such synchronization plates were established for each species. After egg-laying, plates were monitored to ensure 100-200 embryos per plate were laid. *C. inopinata* mixed-sex populations were washed off of plates in M9 buffer at 48 (L3), 66 (L4), and 83 (adult) hours after embryos were laid. *C. elegans* mixed-sex populations were washed off of plates in M9 buffer at 23 (L3), 30 (L4), and 40 (adult) hours after embryos were laid. Before isolating populations, plates were examined to ensure nematodes exhibited morphology consistent with their presumptive developmental stage. After washing, worm concentrations were measured to ensure each tube contained 100 nematodes. Five samples were isolated per species per developmental stage. Nematodes were then resuspended in 250 μl of TRIzol, flash-frozen in liquid nitrogen, and stored at -80°C.

### RNA isolation, library preparation and sequencing

Nematodes in TRIzol were subjected to ten freeze-thaw cycles in liquid nitrogen for tissue disruption. RNA was then isolated with the Qiagen RNeasy kit and resuspended in 15 μl of RNAse-free water. We used 100 ng of total RNA for mRNA extraction and Illumina library preparation using the KAPA mRNA Hyper prep kit (KK8580). Samples were sequenced on the Illumina HiSeq 4000 platform at the University of Oregon (https://gc3f.uoregon.edu/).

### Read processing and transcript abundance inference

Read quality was evaluated with FastQC (with default options) (Andrews 2010). Reads were then demultiplexed with Stacks process_shortreads (with options “-q -c -r --index_null”) (Rochette *et al*. 2019). The barcode for one sample (a *C. inopinata* L4 sample) was not recovered, and this sample could not be included in the analysis. *C. inopinata* (Kanzaki *et al*. 2018) *C. elegans* (Yoshimura *et al*. 2019), *C. briggsae* (Stein *et al*. 2003), *C. nigoni* (Yin *et al*. 2018), and *C. remanei* (Teterina *et al*. 2023) genome assemblies, annotations, mRNA FASTA files, and protein FASTA files were retrieved from WormBase ParaSite (Howe *et al*. 2017). *C. elegans* and *C. inopinata* mRNA FASTA files then were filtered to remove all alternative splice variants except the largest isoform of each gene. These mRNA files were used to generate indices with salmon index (with default options) (Patro *et al*. 2017). *C. inopinata* and *C. elegans* RNA-seq reads were mapped to their respective reference transcriptomes and transcript abundances inferred with salmon quant (with options “-l A -p 8 -- validateMappings –gcBias”) (Patro *et al*. 2017). One *C. elegans* L3 sample revealed a low number of reads that mapped to the reference (Supplemental Table 1). This sample was then excluded from downstream analyses.

All *Caenorhabditis* protein files were filtered to remove alternative splice isoforms (while retaining the largest isoform per gene), and these files were prepared for the OrthoFinder software (with the command “orthofinder -op -S blast -f”) (Emms and Kelly 2019). All pairwise query-database whole-protein searches were performed with blastp (with options “-outfmt 6 -evalue 0.001”) (Camacho *et al*. 2009). Orthologs were identified with OrthoFinder (with options “-S blast -M msa -a 10”) (Emms and Kelly 2019). One-to-one orthologs among *C. elegans* and *C. inopinata* were extracted from the output file “Orthogroups.GeneCount.tsv,” and these defined the 10,718 genes used for downstream analyses of RNA-seq data.

### Differential gene expression and species-stage interaction analyses

Differential gene expression analyses and modeling were performed with DeSeq2 (Love *et al*. 2014), implemented in R (R Core Team). Across all samples, only one gene had a count less than one and was excluded from downstream analyses (leaving 10,717 single-copy orthologs). DeSeq2 fits a generalized linear model of raw gene counts following a negative binomial distribution with a given mean and dispersion for each gene (Love *et al*. 2014); log_2_ fold change coefficients are estimated for each sample (the DeSeq2 function was called with default arguments). DeSeq2 was also used to perform Wald tests for each gene among *C. elegans* and *C. inopinata* at each of the three developmental stages (with the function “results”) (Love *et al*. 2014). Additionally, models including a species-stage interaction term (“∼ Species + Stage + Species:Stage”) were fitted and likelihood ratio tests performed with DeSeq2; in this case, the interaction term gives the estimated difference between the stage effect (across the L4-adult stages) for *C. inopinata* and the stage effect for *C. elegans* (Love *et al*. 2014). *p*-values were corrected for multiple tests with the Holm method (Holm 1979). For PCA and data visualization, counts were regularized log_2_ transformed (function “rlog” with option “blind = FALSE”) (Love *et al*. 2014). Computational workflows, statistical analyses, and data have been deposited in GitHub (https://github.com/gcwoodruff/inopinata_developmental_transcriptomics_2023/)).

### Gene set enrichment analyses

Genes with significant species-stage interactions (Supplemental Table 2) were ranked-ordered by the interaction term and the top 10% (“positive interactions”; Supplemental Table 3) and bottom 10% (“negative interactions”; Supplemental Table 4) of these genes were extracted for ontology analyses. These lists were used as the input for the WormBase “Gene Set Enrichment Analysis” tool (Angeles-Albores *et al*. 2016, 2018) (Figure 4). This reveals enrichment of WormBase Tissue (Lee and Sternberg 2002), WormBase Phenotype (Schindelman *et al*. 2011), and Gene Ontology (Ashburner *et al*. 2000) terms in the input gene list compared to all *C. elegans* genes. The same lists were used as input for the WormExp tool (Yang *et al*. 2016) (Figure 5), which compares the input list with 2,953 gene lists from previous *C. elegans* -omics experiments. KEGG pathway over-representation analyses were performed with the WebGestalt tools (Wang *et al*. 2017).

### Expression of transposon-aligning genes

The TransposonPSI database (https://transposonpsi.sourceforge.net/ ; file “transposon_db.pep”) was used to generate a BLAST database to which the *C. elegans* and *C. inopinata* protein sets were queried with blastp (options “-outfmt 6 -evalue 0.005”) (Camacho *et al*. 2009). *C. inopinata* proteins that aligned to this database (that also did *not* align to any *C. elegans* proteins) were classified as “transposon-aligning” proteins. These were used to compare the two types of genes (those that do and do not align transposons) considering the whole *C. inopinata* gene set in Figure 7 (irrespective of homology with *C. elegans* genes).

The R packages “airway” (Himes *et al*. 2014), “tximport” (Soneson *et al*. 2016), “DESeq2” (Love *et al*. 2014), “PoiClaClu” (Witten 2019), “ggplot2” (Wickham 2016), “ggforce” (Pedersen 2022a), “cowplot” (Wilke 2020), “patchwork” (Pedersen 2022b), “reshape2” (Wickham 2007), “lemon” (Edwards 2020), “GGally” (Schloerke *et al*. 2021), “tidyr” (Wickham *et al*. 2023) were used for this study. Details of computational workflows have been deposited in GitHub (https://github.com/gcwoodruff/inopinata_developmental_transcriptomics_2023/)).

## Results

### Most genes are differentially expressed and exhibit divergent dynamics among species

The length difference between *C. elegans* and *C. inopinata* increases dramatically during the L4-adult transition (Woodruff *et al*. 2018) (Figure 1). To understand the transcriptional basis of body length divergence, we performed RNA-seq on populations of both species at the L3, L4, and adult stages. Differences in reproductive mode among species were accounted for by using *C. elegans fog-2(q71)* animals. This is a *C. elegans*-specific gene encoding an F-box protein implicated in germ line sex determination (Nayak *et al*. 2004). Hermaphrodites homozygous for the *fog-2(q71)* genotype are unable to produce sperm, and this mutation effectively causes *C. elegans* to behave as a female/male species (with obligate outcrossing and a 50:50 sex ratio) (Schedl and Kimble 1988). This allowed both species to harbor mixed-sex populations and facilitated transcriptomic comparisons.

Most genes were differentially expressed at all developmental stages among 10,817 single-copy orthologs. 57%, 55%, and 66% of these genes were differentially expressed between *C. elegans* and *C. inopinata* at the L3, L4, and adult stages respectively (Wald Test Holm-adjusted *p* < 0.05; Figure 2A-C; Supplemental Tables 5-7). To identify genes with divergent dynamics across the key developmental window of interest (the L4-adult transition; Figure 1), we used a model comparison approach to identify genes with significant species-stage interactions with respect to this developmental window (see Methods). This likewise revealed about two-thirds of the single-copy orthologs (67%; 7,204/10,817) exhibit divergent dynamics across this period (Likelihood Ratio Test Holm-adjusted *p* < 0.05; Figure 2D; Supplemental Table 2). Thus, not only are most genes differentially expressed at any given developmental stage, most genes reveal differing dynamics across developmental stages.

**Figure 2.**
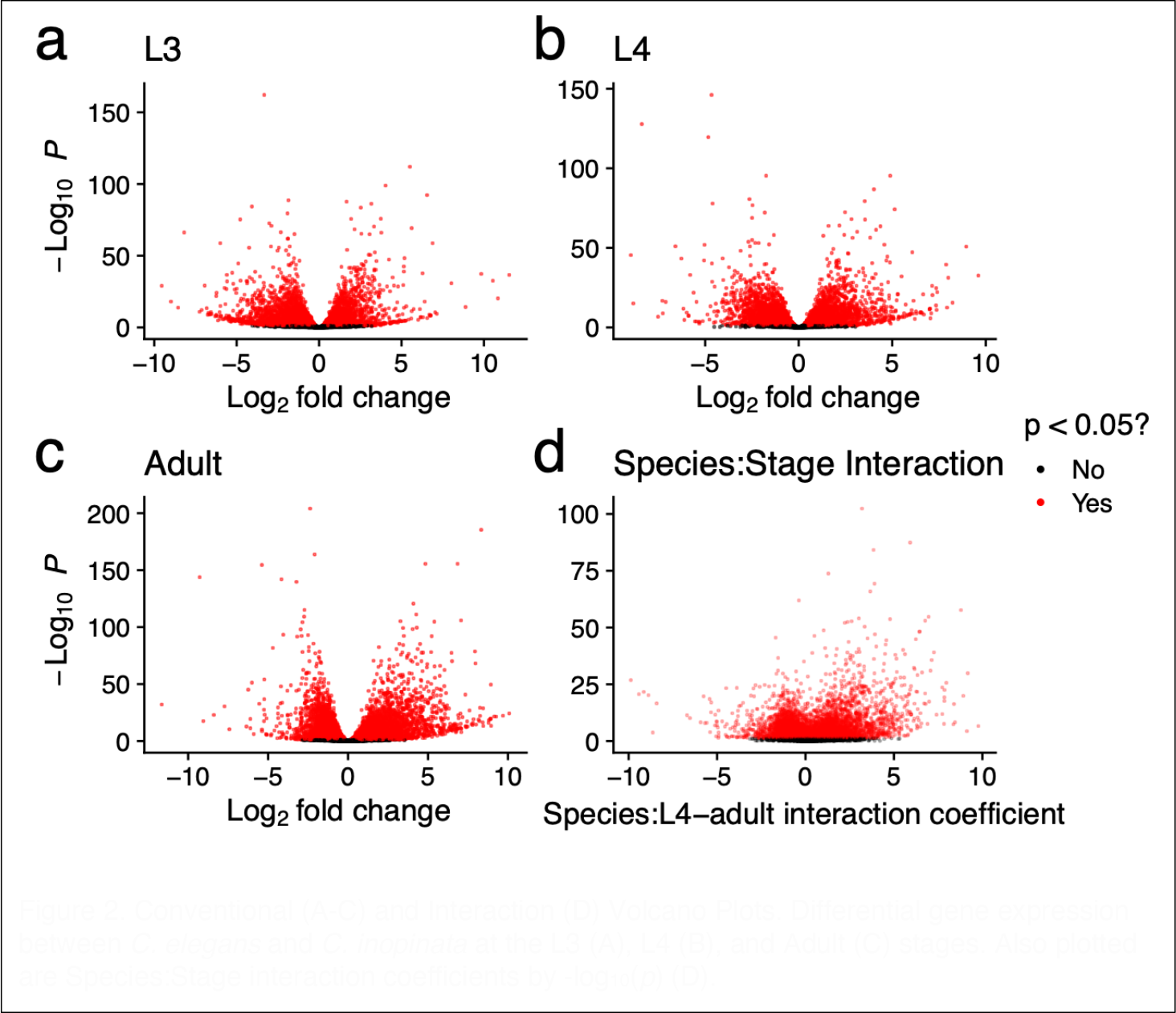
Conventional (A-C) and Interaction (D) Volcano Plots. Differential gene expression between *C. elegans* and *C. inopinata* at the L3 (A), L4 (B), and Adult (C) stages. Also plotted are Species:Stage interaction coefficients by -log_10_(*p*) (D).

### Genes with highly divergent dynamics tend to have behavioral, cuticular, germ line, and stress-response functions

To understand the kinds of genes exhibiting divergent developmental dynamics among species, ontology enrichment analyses were performed. However, as most genes revealed significant species-stage interactions (Figure 2D; Supplemental Table 2), the top ten percent (Supplemental Table 3) and the bottom ten percent (Supplemental Table 4) of these genes as ranked by species-stage interaction coefficient were used for enrichment analyses (Figure 3). These defined the “Positive Interactions” list (720 genes; Supplemental Table 2) and the “Negative Interactions” list (720 genes; Supplemental Table 3). As expected, genes with high species-stage interaction coefficients reveal genes whose expression increase across the L4-adult developmental window in *C. inopinata* but decrease in *C. elegans* (Figure 3A). Genes with low species-stage interaction coefficients reveal the opposite pattern—such genes decrease in expression across this window in *C. inopinata* while increasing in *C. elegans* (Figure 3B). Notably, among the genes with the ten lowest species-stage interaction coefficients, five encode cuticle collagens (Figure 3B).

**Figure 3.**
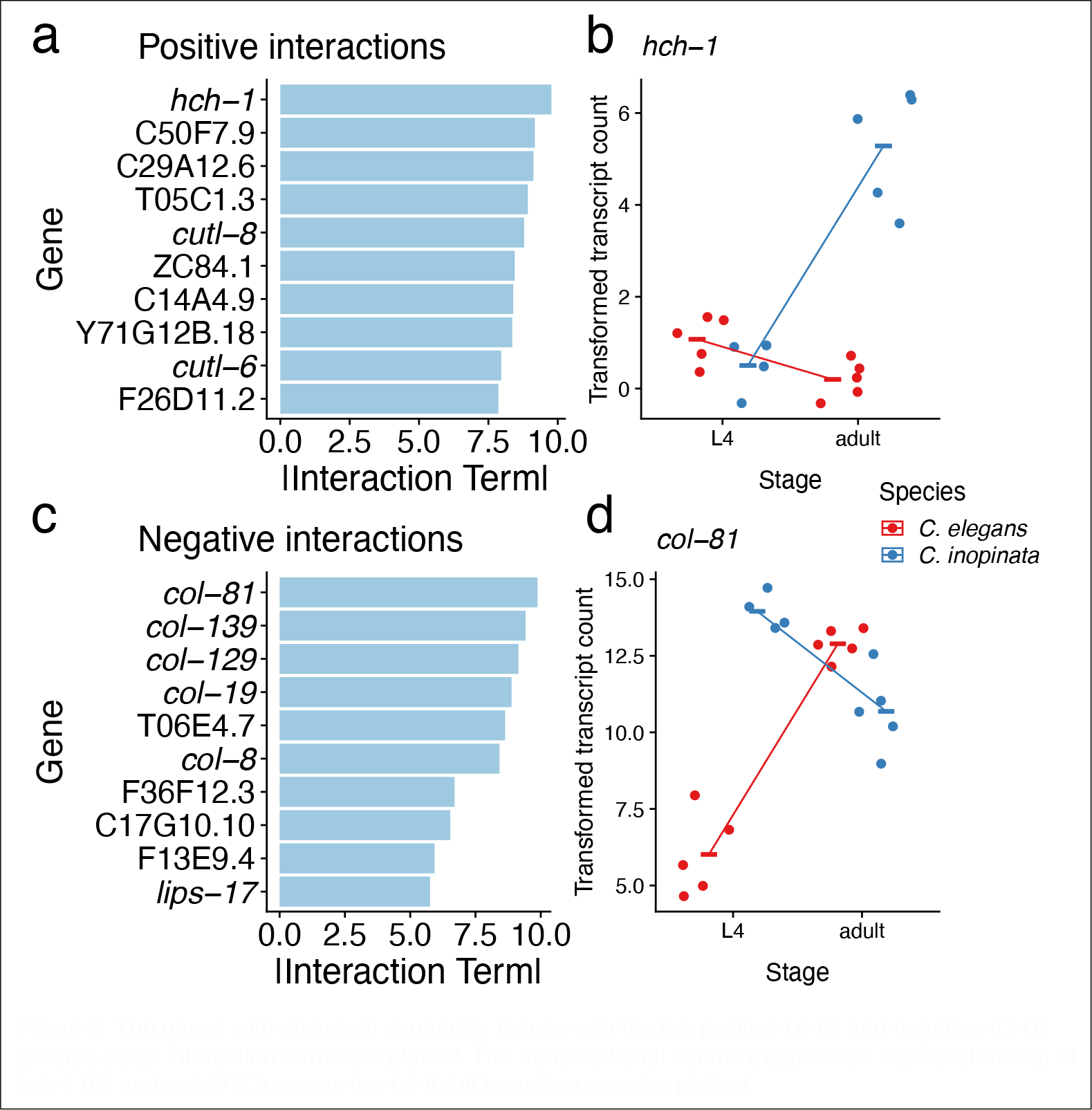
Top genes with divergent dynamics. Genes with the top positive (A-B) and negative (C-D) species-stage interaction terms are plotted. The transcriptional counts (regularized log transformed) of *hch-1* (B) and *col-81* (C) across the L4-Adult transition are also plotted.

Both lists were analyzed with the WormBase Gene Enrichment tool (Angeles-Albores *et al*. 2016, 2018), which compares the frequencies of *C. elegans*-specific tissue and phenotype ontology terms associated with an observed gene list with those expected in the entire *C. elegans* gene set. The genes with positive interactions were enriched for genes expressed in neurons (Figure 4A) and genes with neuronal and behavioral phenotypes upon perturbation (Figure 4B). Additionally, such genes were also enriched for morphological phenotypes such as “dumpy” and “body morphology variant” (Figure 4B). Genes with negative interactions were enriched for germ line, somatic gonad, and early embryonic cell expression (Figure 4C) as well as early-embryo and germ-line phenotypes (Figure 4D). In addition to species-specific tissue and phenotype ontologies, the two lists were also analyzed with the WormExp tool (Yang *et al*. 2016) (Figure 5). This tool compares a gene list from a *C. elegans* genomics study and compares it with a curated collection of such gene lists from previous *C. elegans* experiments; this tool identifies lists with an unexpected degree of overlap. Genes with positive species-stage interactions revealed high overlap with previous *C. elegans* experiments examining stress response (Zarse *et al*. 2012; Bond *et al*. 2014; Burton *et al*. 2017; Delaney *et al*. 2017), small RNAs (Zisoulis *et al*. 2010; Corrêa *et al*. 2010; Padeken *et al*. 2019), and the molting cycle (Hendriks *et al*. 2014) (Figure 5A). Conversely, genes with negative species-stage interactions likewise revealed overlap with studies regarding stress response (Rohlfing *et al*. 2010; Chang *et al*. 2017) and small RNAs (Claycomb *et al*. 2009, 2009; Corrêa *et al*. 2010), as well as the germ line (Claycomb *et al*. 2009; Boyd *et al*. 2010; Kershner and Kimble 2010; Greer *et al*. 2010; Gracida and Eckmann 2013), epidermal collagens (Rohlfing *et al*. 2010), and neuromuscular lamins (González-Aguilera *et al*. 2014). A KEGG pathway analysis of the genes with negative interactions revealed over-representation of the categories “Base excision repair,” “Homologous recombination,” and “DNA replication,” consistent with germ line and early embryo functions (genes with positive interactions revealed no significant over-represented KEGG categories). Thus, although these analyses revealed enriched functions related to obvious features of phenotypic divergence such as body morphology (Figure 4B; Figure 5B) and developmental timing (Figure 5A), they also revealed surprising sets of genes connected to neuronal, germ line, and stress-response roles.

**Figure 4.**
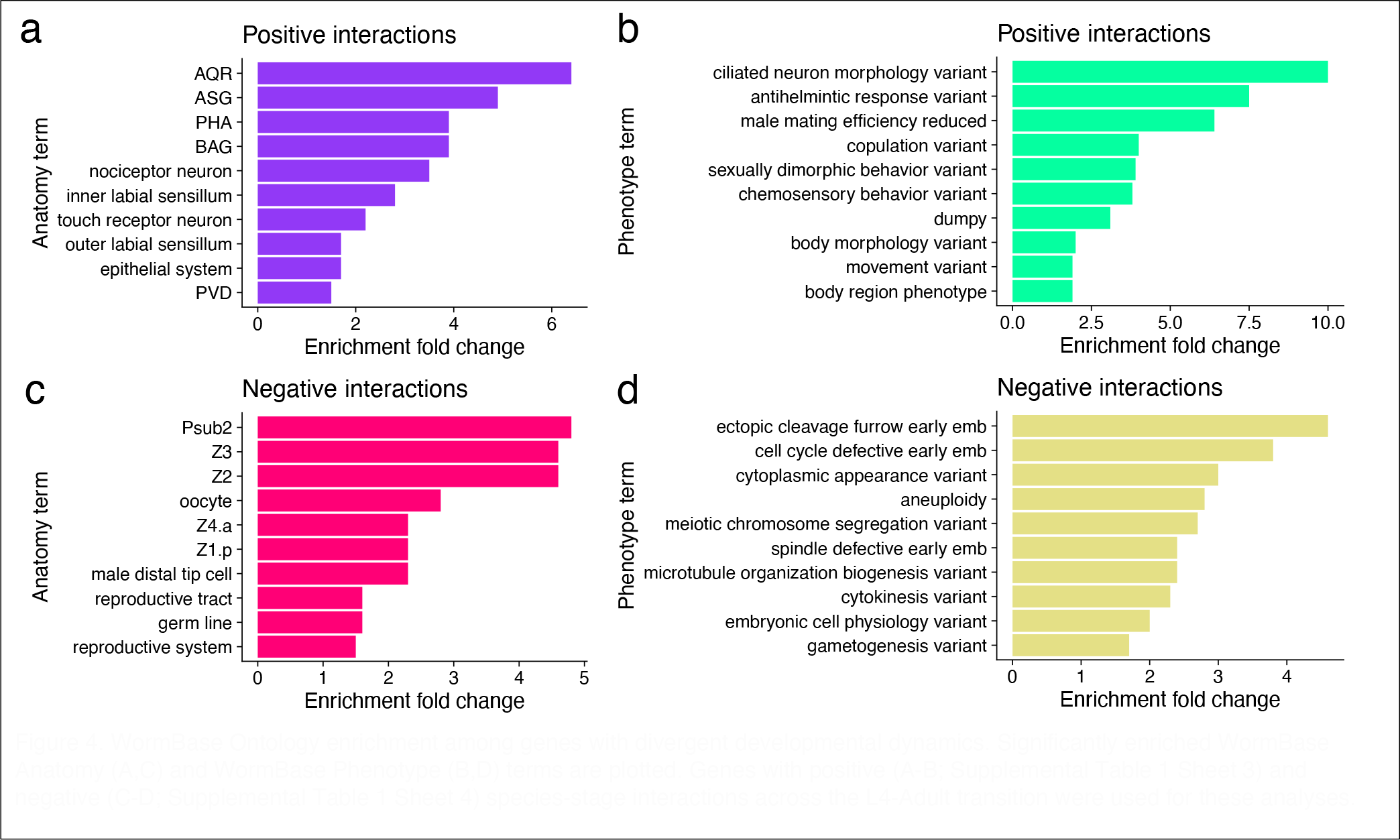
WormBase Ontology enrichment among genes with divergent developmental dynamics. Significantly enriched WormBase Anatomy (A,C) and WormBase Phenotype (B,D) terms are plotted. Genes with positive (A-B; Supplemental Table 1 Sheet 3) and negative (C-D; Supplemental Table 1 Sheet 4) species-stage interactions across the L4-Adult transition were used for these analyses.

**Figure 5.**
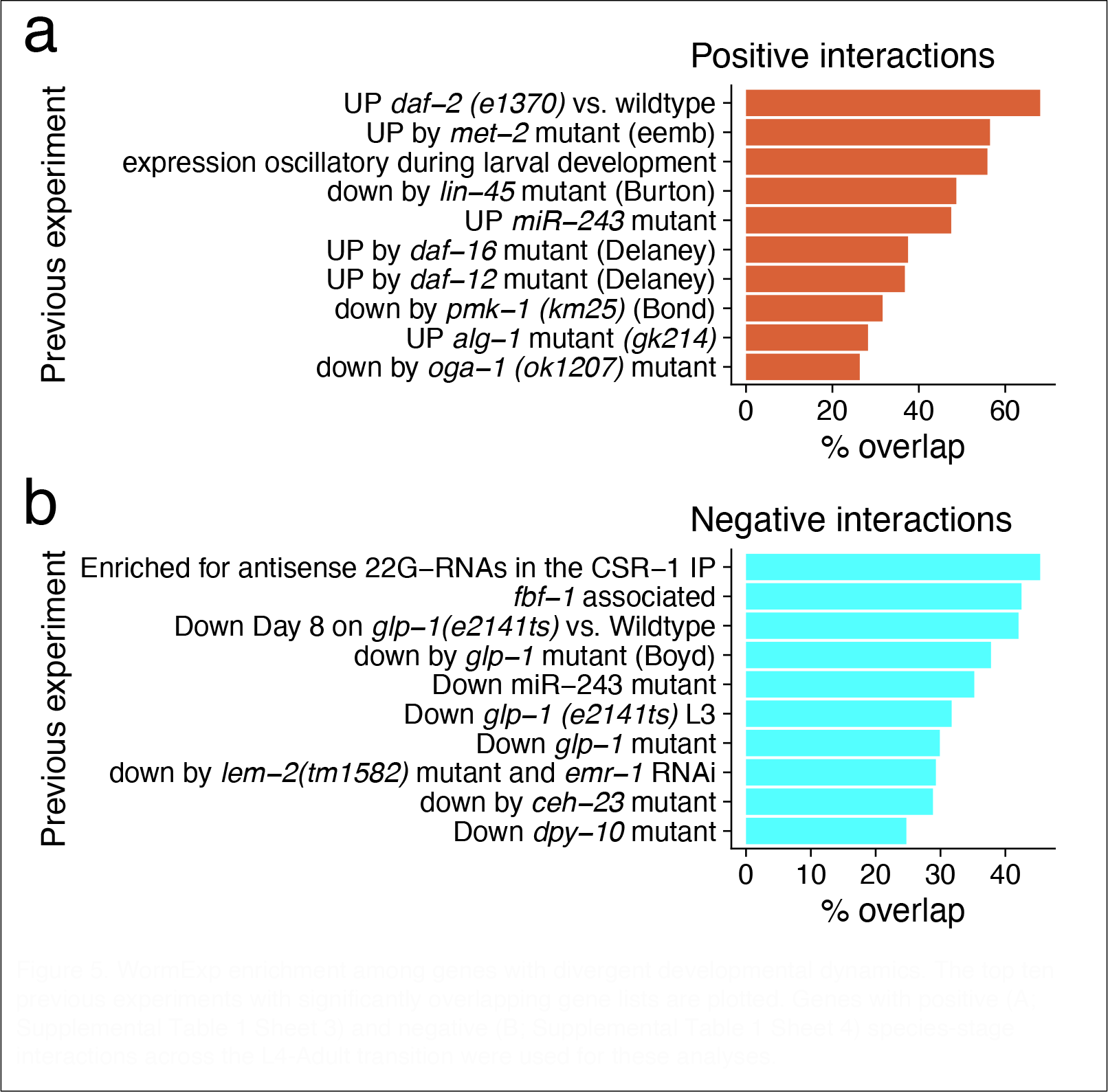
WormExp enrichment among genes with divergent developmental dynamics. The top ten previous experiments with significantly overlapping gene lists are plotted. Genes with positive (A; Supplemental Table 1 Sheet 3) and negative (B; Supplemental Table 1 Sheet 4) species-stage interactions across the L4-Adult transition were used for these analyses.

### Genes associated with TGF-β signaling reveal unexpected patterns of transcriptional divergence

In *C. elegans*, body size is regulated by canonical TGF-β signaling (Savage-Dunn and Padgett 2017). Many genes in this pathway impact body size when perturbed (Savage-Dunn and Padgett 2017). For instance, loss-of-function mutations in the extracellular proteins LON-1 (Morita *et al*. 2002) and LON-2 (Gumienny *et al*. 2007) promote increased body size. Additionally, proteins such as the TGF-β ligand DBL-1 reveal dose-dependent effects on body size—loss-of-function mutants are small whereas overexpression mutants are long (Suzuki *et al*. 1999; Morita *et al*. 1999). Background knowledge regarding the phenotypic effects of mutations in this pathway can thus inform hypotheses regarding the developmental basis of body size evolution. For instance, *C. inopinata* may be long because it has increased DBL-1 expression and/or decreased LON-1 expression. To address these and other possibilities, we examined differential patterns of twenty-six genes associated with TGF-β signaling that were identified in a previous review (Gumienny and Savage-Dunn 2013) (Figure 6).

**Figure 6.**
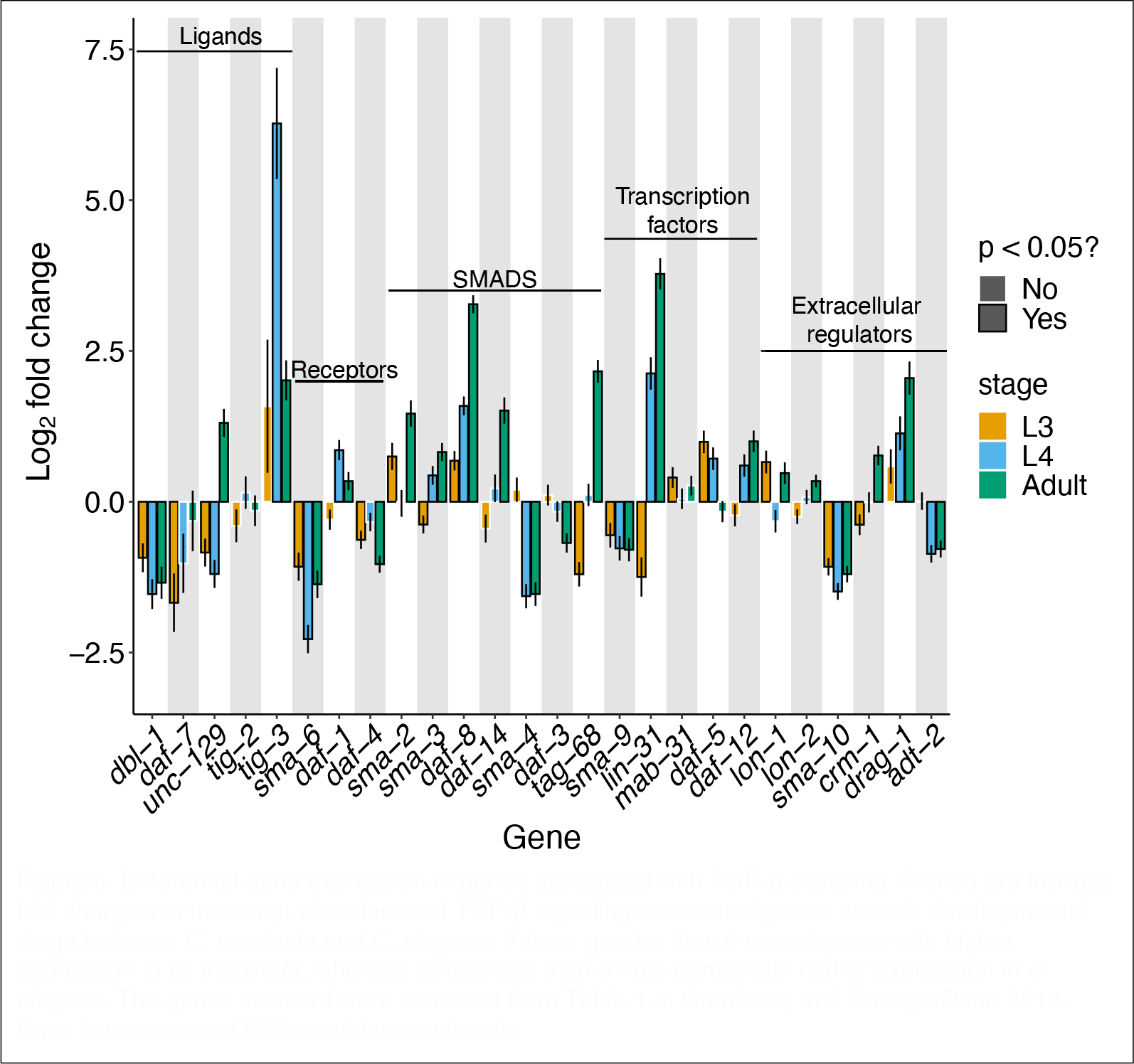
Differential gene expression in genes associated with TGF-β signaling. Plotted are the log_2_ fold changes in transcript abundance of TGF-β signaling-associated genes at each developmental stage between *C. inopinata* and *C. elegans*. Values greater than 0 reveal genes with higher expression in *C. inopinata*, whereas values less than 0 note genes with higher expression in *C. elegans*. The genes included were extracted from Table 1 of Gumienny and Savage-Dunn 2013. Error bars represent 95% confidence intervals.

Like most genes, there is extensive differential gene expression between *C. elegans* and *C. inopinata* across TGF-β pathway genes (Figure 6). However, their differential expression is idiosyncratic (i.e., varied in direction; Figure 6) and often discordant with the increased body size of *C. inopinata*. For instance, *dbl-1* reveals lower expression in *C. inopinata* at all developmental stages compared to *C. elegans* (Figure 6), contrary to expectations from the literature (Suzuki *et al*. 1999). Additionally, although *lon-1* and *lon-2* might be expected to have lower expression in the elongated *C. inopinata* (Morita *et al*. 2002; Gumienny *et al*. 2007), the difference in expression compared to *C. elegans* is negligible or even greater (Figure 6). Yet, a handful of these genes reveal an increase in differential expression across the L4-adult transition (*tig-3, daf-8, lin-31*, and *drag-1*; Figure 6), although these genes are not reported to control body size in *C. elegans*.

Patterns of differential expression of TGF-β signaling genes then do not straightforwardly align with the hypothesis that this pathway drives body size evolution in *C. inopinata*. But it is possible that body size mutants in *C. elegans* harbor a transcriptomic signature that is mirrored in *C. inopinata*. To address this, the top decile of genes with significant species-stage interactions (used in gene enrichment analyses; Fig. 4-5) were compared with gene lists from previous studies measuring transcriptomic changes in *C. elegans* TGF-β pathway mutants (Liang *et al*. 2007; Roberts *et al*. 2010; Lakdawala *et al*. 2019). The top decile of differentially expressed genes across the L3, L4, and adult stages (Fig. 2) were also compared with the gene lists from these studies. No significant overlap was detected between the most differentially expressed (Fig. 2A-C) or dynamically divergent (Fig. 2D) genes identified here and those associated with *C. elegans* TGF-β pathway mutants (Hypergeometric test Holm-adjusted *p* > 0.05). However, the gene list of one study was dominated by collagen genes (15/18 significant differentially expressed genes encoded collagen proteins) (Lakdawala *et al*. 2019), reminiscent of genes with negative species-stage interactions (Fig. 3B). Thus, while the transcriptome of *C. inopinata* has not diverged in a manner that strictly overlaps patterns seen in TGF-β pathway mutants, many cuticle collagens are differentially expressed in such mutants and exhibit divergent developmental dynamics across species.

### Genes that align to transposons exhibit lower expression

*C. inopinata* does not only harbor a unique body size and morphology—it also harbors an unusual repetitive genomic landscape (Kanzaki *et al*. 2018; Woodruff and Teterina 2020). *C. inopinata* maintains many open reading frames encoding proteins related to transposable elements (Woodruff and Teterina 2020). To understand the biological activity of these genes, we compared transcriptional abundances of these transposon-aligning genes with those that do not align to transposons (Figure 7). Across all developmental stages observed, transposon-aligning genes harbor far lower expression than genes that do no align to transposons (Fig. 7; 57-60% reduction in transformed transcript count; Cohen’s *d* effect size = -0.82 - -0.73; Wilcoxon rank-sum test *p* < 0.001). Despite this, there are transposon-aligning genes with high transcriptional abundance (Fig. 7), suggesting these open reading frames maintain some degree of biological activity.

**Figure 7.**
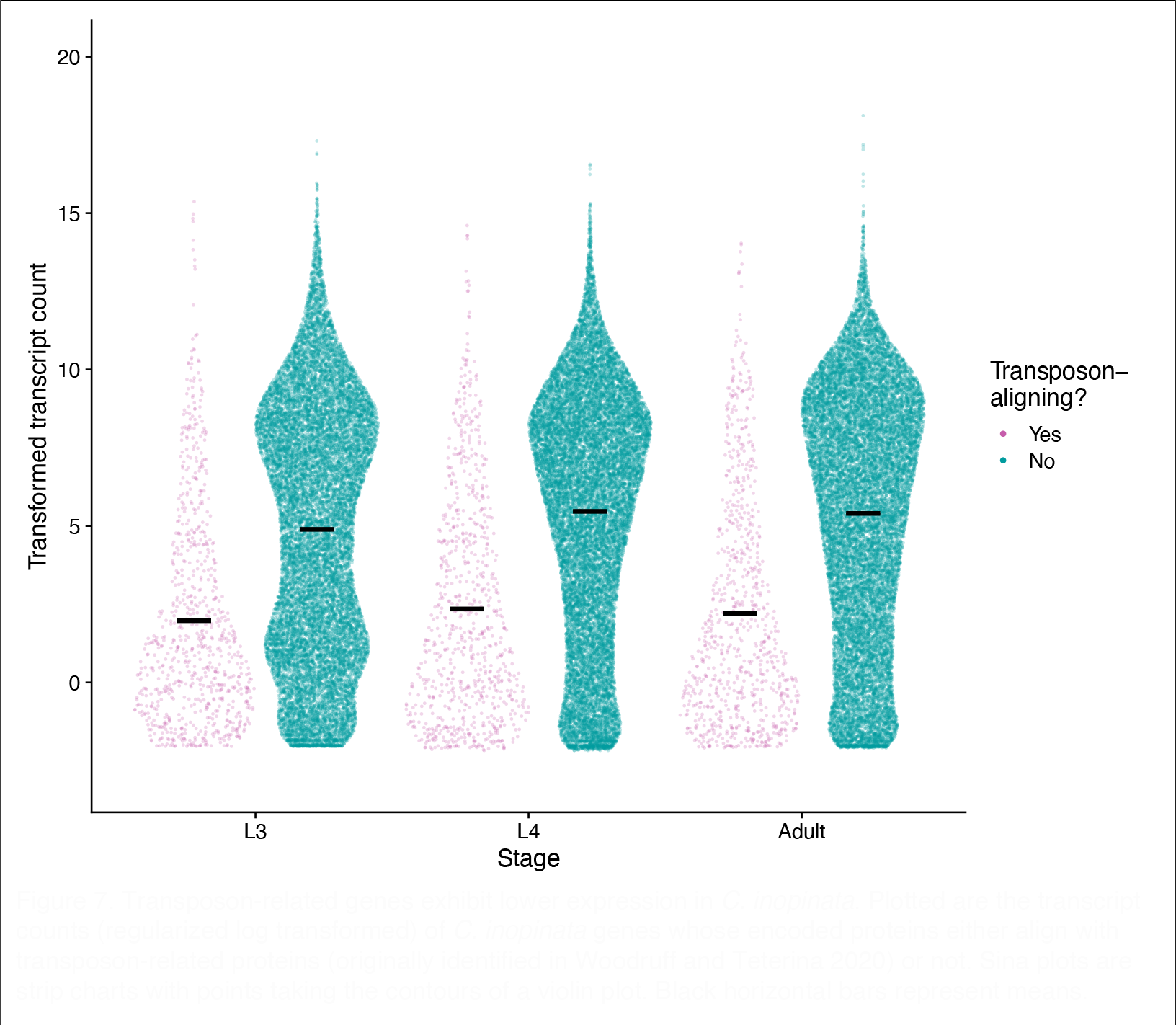
Transposon-related genes exhibit lower expression in *C. inopinata*. Plotted are the transcript counts (regularized log transformed) of *C. inopinata* genes whose encoded proteins either align with transposon-related proteins (originally identified in Woodruff and Teterina 2020) or not. Sina plots are strip charts with points taking the contours of a violin plot. Black horizontal bars represent means.

## Discussion

### Widespread transcriptional divergence across postembryonic development

Over half of about 11,000 single copy orthologs are differentially expressed between *C. elegans* and *C. inopinata* at all developmental stages considered. This is surprising because these species are closely related and harbor similar developmental patterns (Woodruff *et al*. 2018; Kanzaki *et al*. 2018). No differences in the number of somatic cells between *C. elegans* and *C. inopinata* adults could be detected in a previous study (Woodruff *et al*. 2018), suggestive of a highly conserved cell lineage among the two species. Thus, we might potentially expect highly conserved patterns of gene expression to co-occur with these developmental similarities. One potential explanation for these differences could be the use of a *fog-2* mutation in our *C. elegans* populations; this is an unlikely driver of apparent rampant gene expression divergence because this gene’s function appears to be limited to hermaphrodite germ-line sex determination and has no other clear impacts on fitness (Schedl and Kimble 1988). A more likely explanation for the expression divergence is the morphological and ecological divergence among these species. Not only is *C. inopinata* much longer than *C. elegan*s— it also thrives in a markedly different natural environment (fresh *F. septica* figs instead of rotting plants)(Kanzaki *et al*. 2018; Woodruff and Phillips 2018). Its radically divergent transcriptome may then reflect its divergent morphology and ecology, and *C. inopinata* in particular may require the ubiquitous tuning of gene expression to shape its needs as a fig nematode.

Another possible explanation is that such divergent patterns of gene expression are common among closely-related nematode species. In other words, developmental system drift may explain these differing transcriptomic dynamics (True and Haag 2001). The extent of differential gene expression across postembryonic development in *Caenorhabditis* nematodes is not entirely obvious.

Unexpectedly divergent expression across embryogenesis has been reported in *C. elegans* and *C. briggsae* (Yanai and Hunter 2009). Alternatively, it has been reported that gene expression across postembryonic development is largely conserved between *C. elegans* and *C. briggsae* (Grün *et al*. 2014; Lu *et al*. 2020). It has also been shown that transcriptional patterns are more likely to be conserved during ventral enclosure when compared to other embryonic stages (Levin *et al*. 2012). Additionally, hermaphroditic species tend to have less complex and less sex-biased transcriptomes than gonochoristic species (when considering adults) (Thomas *et al*. 2012), and a transgenic reporter construct survey with eight genes and four *Caenorhabditis* species revealed widespread spatial variation in gene expression (Barrière and Ruvinsky 2014). Notably, over half of the genes examined were found to be differentially expressed between *C. briggsae* and *C. nigoni* nematodes of the same sex (Sánchez-Ramírez *et al*. 2021). As these species are far more closely related to each other than *C. inopinata* and *C. elegans* (Sloat *et al*. 2022), this lends support to the view that developmental system drift in transcriptomes is common in this group. Regardless, future studies that capture a larger phylogenetic sample as well as a range of postembryonic stages will be required to disentangle these possibilities.

### Transcriptional divergence and body size evolution

We were able to identify transcriptional patterns of many orthologs connected to TGF-β signaling. This pathway influences body size in *C. elegans*, and numerous mutants associated with this pathway exhibit small or elongated bodies. Surprisingly, our results revealed idiosyncratic patterns of differential gene expression that do not align with simple models of TGF-β signaling (Fig. 6). For instance, in *C. elegans*, high levels of *dbl-1* transcription promote repression of the downstream target gene *lon-1*, leading to body size increases (Morita *et al*. 2002). Thus, a natural hypothesis would be that *C. inopinata* is long due to increased *dbl-1* (and decreased *lon-1*) expression. Neither of these patterns were observed; *dbl-1* is *lower* in expression while *lon-1* exhibits negligible differences in gene expression (Fig. 6). Thus, it is unlikely that body size is driven by transcriptional evolution of *dbl-1, lon-1*, or other TGF-β signaling genes in a manner concordant with such simple hypotheses derived from *C. elegans* developmental genetics. This is consistent with the observation that *C. inopinata* does not harbor differences in hypodermal endoreplication compared to *C. elegans* (Woodruff *et al*. 2018); TGF-β signaling has been proposed to regulate body size through this mechanism (Lozano *et al*. 2006). However, gene functions have been shown to evolve in *Caenorhabditis* nematodes (Beadell *et al*. 2011; Verster *et al*. 2014), and it is entirely possible that the roles of TGF-β pathways have changed in *C. inopinata* (which can then resolve these unexpected transcriptional patterns). Future studies involving the perturbation of these genes’ activities in *C. inopinata* will be required to interrogate this possibility.

Notably, cuticle collagens were common among genes with negative species-stage interactions (Fig. 3). *C. elegans* harbors over a hundred such genes (Teuscher *et al*. 2019), and a number of genes with morphological mutant phenotypes encode such collagens (Johnstone 2000) (although most collagen genes have no described phenotypes). As these genes encode core components of the extracellular matrix constituting the exoskeleton, it is unsurprising that some of these genes regulate body morphology. Moreover, collagen genes have been shown to be regulated by TGF-β signaling (Madaan *et al*. 2018, 2020), and this pathway may regulate body morphology in part through controlling the expression of such exoskeletal factors. Thus, it is possible that these divergent transcriptional dynamics in collagen genes may promote the evolution of elongated body size in *C. inopinata*.

### Stress, behavior, small RNAs, and the germ line

Enrichment analysis of divergently dynamic genes generated results concordant with *C. inopinata*’s divergent body size (“body morphology variant” and “dumpy” WormBase Phenotypes; Fig. 4) and developmental rate (“expression oscillatory during larval development” WormExp experiment (Hendriks *et al*. 2014); Fig. 5). However, enrichment analyses also detected a range of unexpected biological phenomena associated with divergent developmental transcriptional dynamics. For instance, overlap was found between divergently dynamic genes and genes exhibiting differential expression in a number of experiments related to stress-response (including genes such as *daf-2* (Zarse *et al*. 2012), *daf-16* (Delaney *et al*. 2017), and *daf-12* (Delaney *et al*. 2017); Fig. 5a). Additionally, genes connected to neurons (Fig. 4a) and behavioral phenotypes (Fig. 4b) were also enriched. Some of these behavioral phenotypes (such as “copulation” and “male mating efficiency”) are likely due to the difference in reproductive mode between species. *C. elegans* is a self-fertile hermaphrodite harboring low male frequencies in natural populations (Cutter *et al*. 2019); *C. inopinata* is an obligate outcrosser (Kanzaki *et al*. 2018). However, we speculate these other differences result from these species’ divergent natural ecological contexts. *C. elegans* thrives in rotting plant material (Frézal and Félix 2015) and grows readily in laboratory conditions. *C. inopinata* thrives in fresh figs (Kanzaki *et al*. 2018; Woodruff and Phillips 2018) and has low fecundity in laboratory conditions (Woodruff *et al*. 2019). Thus, its behavioral and stress-response regimes are likely to be tuned to a radically different natural context, thus driving patterns of divergently developmentally dynamic expression in the genes underlying these biological functions. Consistent with this, the stress-resistant dauer stage has diverged in *C. inopinata*, exhibiting an apparent loss of radial constriction and a far lower prevalence in laboratory conditions than *C. elegans* (Hammerschmith *et al*. 2022). Thus, while it is possible there may be some co-option of behavioral and stress genes in the divergent growth and developmental processes of *C. inopinata*, it is more likely that these traits themselves have diverged in this lineage.

Germ line genes were also detected in both enrichment analyses performed (Fig. 4c-d, Fig. 5b). This is also likely due to *C. inopinata*’s divergent environmental context and low fecundity in laboratory conditions. A previous study revealed the adult female gonad of *C. inopinata* is much smaller and holds far fewer germ cells than that of *C. elegans* (Woodruff *et al*. 2018). Indeed, if somatic and germ cells are included, *C. elegans* has *more* cells than *C. inopinata* despite its smaller body size (Woodruff *et al*. 2018). Additionally, these enrichment analyses were performed on samples undergoing maturation— if patterns of oogenesis and early embryogenesis are divergent (which has been observed in *Caenorhabditis* (Yanai and Hunter 2009; Levin *et al*. 2012; Farhadifar *et al*. 2015)), then it would be unsurprising to see such divergent dynamics of germ line genes in our samples. Connected to this, genes associated with small RNA biology were also detected in enrichment analyses (such as *csr-1* (Claycomb *et al*. 2009), *alg-1* (Zisoulis *et al*. 2010), *mir-243* (Corrêa *et al*. 2010); Fig. 5). This may simply reflect the potential germ line divergence described above. The germ line harbors an array of tissue-specific granules that contain small RNAs (Sundby *et al*. 2021), and germ line divergence may entail the evolution of germ granules. However, small RNAs (particularly piRNAs) are known to regulate transposable elements (Tóth *et al*. 2016), and the genome of *C. inopinata* has evolved a highly repetitive (Kanzaki *et al*. 2018) and surprisingly uniform (Woodruff and Teterina 2020) landscape of such elements. Moreover, *C. inopinata* has lost the key small RNA regulators *ergo-1, eri-9*, and *eri-6/7* (Kanzaki *et al*. 2018). The divergent dynamics of small RNA genes may also then be connected to *C. inopinata*’s exceptionally transposable element-rich genome and the loss of conserved small RNA machinery.

### Transposable elements

In addition to having a transposon-rich genome in general, *C. inopinata* has many open reading frames (ORFs) that encode transposon-related proteins (such as integrases, polymerases, ribonucleases, etc.) (Woodruff and Teterina 2020). Transposons are expected to be deleterious to the host (Wicker *et al*. 2007), and as a consequence, myriad defenses have evolved to silence such elements and inhibit their activity (Buchon and Vaury 2006; Tóth *et al*. 2016). Here, we showed that these transposon-aligning open reading frames reveal a 50% reduction in mean transformed transcript count compared to genes that do not align to transposons (Fig. 7). This is consistent with these ORFs being deleterious and with host inhibition of their activity. However, many of these genes are highly expressed (Fig. 7). This suggests that many of these genes are active transposable elements or otherwise harbor activity that is biologically-relevant to the host. Disentangling these possibilities will require whole-genome sequencing of either multiple *C. inopinata* populations (which has been reported to be in progress (Kawahara *et al*. 2023)) or longitudinal studies of single populations to track the transposition of active elements. Indeed, a recent study showed that *C. inopinata*-specific transposable element insertions are associated with changes in gene expression across species (Kawahara *et al*. 2023). Beyond this, it remains possible that transposon-associated ORFs have been co-opted for host functions (Jangam *et al*. 2017; Singh and Bhalla 2020), and future studies will be required to address this possibility.

### Findings in light of Kawahara et al. GBE 2023

Recently, a study performing a very similar set of experiments was published (Kawahara *et al*. 2023). How do our results compare? For instance, under principal components analysis, the first two principal components of both studies separate samples of differing stages and species (compare Suppl. Fig 1 of Kawahara *et al*. 2023 with Suppl. Fig. 1 here). Moreover, they found that 64-71% of single-copy orthologs were differentially expressed across species, even greater than our observations (55-66%). Thus, both studies suggest body size evolution is accompanied by widespread gene expression divergence. Additionally, Kawahara *et al*. reported notable divergence in collagen gene expression, as well as idiosyncratic expression across the TGF-β signaling pathway, consistent with our findings. Additionally, their detailed analyses of transposable element insertion-impacts on ortholog expression are consistent with our findings that transposon-aligning genes are expressed. Thus, our second report confirms these broad conclusions are robust. That said, there are more specific, quantitative differences between these reports. For instance, Kawahara *et al*. detected no differential expression in *dbl-1* (their Suppl. Fig. 6), whereas we found this gene is under-expressed in *C. inopinata* at all developmental stages (Fig. 6). While it is unclear exactly what is driving these discordant findings, it is worth noting there were a number of biological differences among the studies that may explain specific differences in transcript abundances. For instance, in our study, all animals were grown at 25°C on *E. coli* OP50-1. In Kawahara *et al*. 2023, *C. inopinata* was grown on *E. coli* strain HT115 (DE3) at 27°C, while *C. elegans* was grown at 24.5°C. Additionally, we used *C. elegans fog-2 (q71)* to account for reproductive mode, whereas Kawahara *et al*. 2023 used *fem-3 (hc17)*. Indeed, our populations included males, whereas Kawahara *et al*. 2023 only examined females. Thus, the environments, sexual compositions, and genetic backgrounds differed across these studies; this is likely to explain such specific differences in transcription. It is thus all the more striking that the broad conclusions of these studies are robust to such differences in experimental design.

### Concluding thoughts

Here, we found widespread transcriptional divergence across the larval-to-adult transition between *C. elegans* and *C. inopinata*. While the extent of developmental system drift in transcript abundance in this group is uncertain, some fraction of these transcriptionally divergent genes must be implicated in the evolution of increased body length in *C. inopinata*. Genes with divergent dynamics included those encoding collagens, those with body morphology size phenotypes in *C. elegans*, and those connected to TGF-β signaling. This work then reveals multiple specific hypothetical drivers of body size in this group and sets the stage for future laboratory experiments interrogating the developmental and genetic bases of body size evolution.

## Data availability

FASTQ files have been submitted to the NCBI Sequence Read Archive (SRA; http://www.ncbi.nlm.nih.gov/sra) under the BioProject ID PRJNA1031217. Sample metadata can be found in supplemental_tables.xls Sheet 1. All other data and code affiliated with this work have been deposited in Github (https://github.com/gcwoodruff/inopinata_developmental_transcriptomics_2023).

## Acknowledgements

We thank the University of Oregon Genomics and Cell Characterization Core Facility (GC3F) for assistance with Illumina sequencing. This work also benefited from access to the University of Oregon high performance computer, Talapas. Some of the computing for this project was performed at the OU Supercomputing Center for Education & Research (OSCER) at the University of Oklahoma (OU). We also thank WormBase and the curators of WormExp (Wentao Yang et al.).

## Author contributions

GCW and PCP devised and designed the project. GCW and EJ reared nematodes and prepared RNA samples. JHW prepared libraries for sequencing. GCW performed bioinformatic analyses and wrote the first draft of the paper. GCW, EJ, JHW, and PCP revised and prepared the final manuscript.

## Author notes

The authors declare no conflict of interest.

